# Antibacterial activity of *Xenopsylla cheopis* attacins against *Yersinia pestis*

**DOI:** 10.1101/2023.08.26.554949

**Authors:** Basil Mathew, Kari L Aoyagi, Mark A. Fisher

## Abstract

Antimicrobial peptide resistance has been proposed to play a major role in the flea-borne transmission of *Yersinia pestis*. However, the antimicrobial peptide response in fleas and their interaction with *Y. pestis* is largely unknown. Attacins are one of the most abundantly expressed antimicrobial peptides within the first hours after *Y. pestis* infection of *Xenopsylla cheopis*, a major vector of plague. In this study, we report the cloning, expression, and purification of two *X. cheopis* attacin peptides and describe their interactions with and antimicrobial activities against *Y. pestis*. These flea attacins were shown to bind lipopolysaccharides and have potent activity against *Y. pestis*, however the mechanism of killing does not involve extensive membrane damage. Treatment with attacins rapidly inhibits *Y. pestis* colony formation and results in oxidative stress, yet live-cell imaging revealed that bacteria continue to grow and divide for several hours in the presence of attacins before undergoing morphological changes and subsequent lysis. This data provides insights into an early battle between vector and pathogen that may impact transmission of one of the most virulent diseases known to man.

## Introduction

Transmission of *Yersinia pestis*, the causative agent of plague in the wild is mediated by fleas, a group of hematophagous insects belonging to the order Siphonaptera (*1, 2*). Antimicrobial peptide resistance is important for the flea-borne transmission of *Y. pestis* (*3, 4*). The oriental rat flea, *Xenopsylla cheopis*, is considered to be one of the most efficient vectors of plague (*1, 5, 6*). Despite being a major vector of one of the most virulent human pathogens, very little is known about the interactions between *Y. pestis* and flea antimicrobial peptides (AMPs) (*7, 8*). Recently published cat flea genome (*9*) and transcriptomic studies (*10-12*) have provided valuable insights into the range of potential immune responses in fleas, particularly regarding AMP responses. Using these resources, we recently characterized a cecropin class AMP named cheopin from *X. cheopis*, and described its interactions with *Y. pestis* (*7*). To date, this is the only characterized AMP from the insect order Siphonaptera.

Unlike many other *Enterobacterales* species, *Y. pestis* has a rough type lipopolysaccharide (LPS) which contains lipid A and core oligosaccharides but lacks O-antigen polysaccharides (*13*). Modifications of LPS at the lipid A head groups, lipid A fatty acyl chains, and core oligosaccharides have all been reported to confer bacterial resistance against cationic AMPs (*14*). The replacement of lipid A phosphate groups with 4-amino-4-deoxy-L-arabinose (Ara4N) is critical for resistance against the *X. cheopis* AMP cheopin (*7*) and multiple other AMPs (*13, 15, 16*). The activity of cecropin A on *Y. pestis* is affected by the lipid A acylation status as demonstrated by a strain producing primarily hexaacylated lipid A being more resistant to cecropin A than a mutant expressing only tetraacylated lipid A (*17*). Variations in the core oligosaccharide may also increase resistance against polymyxin B as shown in a *Y. pestis phoP* mutant strain (*18*). In vitro screening of *Y. pestis* mutant libraries against AMPs such as polymyxin B and host-defense peptides from insects and mammals indicated potential LPS-independent mechanisms of AMP resistance (*3, 19*). Absence of a universal mechanism of resistance was further evidenced by the fact that *Y. pestis* mutants that were resistant to polymyxin B were susceptible to protamine or cathelicidin, and vice versa (*19*). These studies suggest that different resistance mechanisms may have evolved against structurally distinct classes of AMPs. Therefore, characterization of different families of flea AMPs is necessary to improve our understating of flea-borne transmission of plague.

In a recent study, Bland et al. analyzed transcriptomic changes in the *X. cheopis* digestive tract in response to *Y. pestis* infection and reported that multiple classes of AMPs are expressed in response to *Y. pestis* infection. Two transcripts encoding attacin-class AMPs were significantly induced in response to *Y. pestis* infection (*10*). Attacins are large (∼20 kDa) linear antimicrobial peptides of insect origin with antibacterial activity against some Gram-negative bacteria (*20, 21*). Although the exact mechanisms of action of attacins are not yet completely understood, a proposed mechanism involves interaction with LPS (*22, 23*) and subsequent deregulation of outer membrane homeostasis by inducing expression of multiple outer membrane proteins such as OmpA, OmpF, and OmpC (*23, 24*). Though attacins are known to interact with bacterial outer membranes and LPS through electrostatic interactions, they can retain activity against bacteria with Ara4N-modified LPS that are resistant to other AMPs such as polymyxin B (*23*). Although *X. cheopis* attacins are significantly induced in response to *Y. pestis* infection (*10*), their physiological relevance, antibacterial activities, and interactions with *Y. pestis* are unknown.

In this study we describe the antimicrobial activity and mechanisms of action of two *X. cheopis* attacins, attacin-1 and attacin-2 (Fig S1), which are induced 60- and 35-fold respectively early in *Y. pestis* infection of fleas (*10*). Our results show that these attacins have potent activity that appears targeted against *Y. pestis* and closely related species. Investigations into the mechanisms of action indicate that attacins elicit their bactericidal activity through a mechanism where bacteria continue to grow and divide for several hours before undergoing morphological changes followed by lysis.

## Results

### X. cheopis attacins exhibit potent activity against Y. pestis

The attacins, particularly attacin-1, showed substantial activity against *Y. pestis*, with minimal inhibitory concentrations (MIC) of 1.8 μM (32 μg/mL) and 7.3 μM (128 μg/mL) for attacin-1 and attacin-2, respectively (Table 1). However, the peptides showed no activity against Gram-positive and Gram-negative quality control strains at the highest concentration tested, with MICs ≥ 14.4μM (≥256 μg/mL) for all species. Deletion of the *arn* operon, which encodes for amino arabinose (Ara4N) modification of LPS that is responsible for resistance to multiple cationic AMPs (*3, 7*), only reduced the attacin-1 MIC by a single two-fold dilution. In contrast, the absence of Ara4N-modified LPS resulted in an 8-fold decrease in the MIC of attacin-2. The expression of *Yersinia pseudotuberculosis* O-antigen oligosaccharides in *Y. pestis* did not influence attacin MICs, which is consistent with the fact that a *Y. pseudotuberculosis* clinical isolate had the same attacin-1 MIC as *Y. pestis*. Because attacins exhibit activity against *Y. pestis* and *Y. pseudotuberculosis*, which are both resistant to polymyxins, we tested their activity against a broader panel of polymyxin B resistant strains (Table S2). The attacins showed no discernible activity against these strains, which suggests that *X. cheopis* attacins possess selective activity against *Yersinia* spp., particularly *Y. pestis*.

**Table 1.**
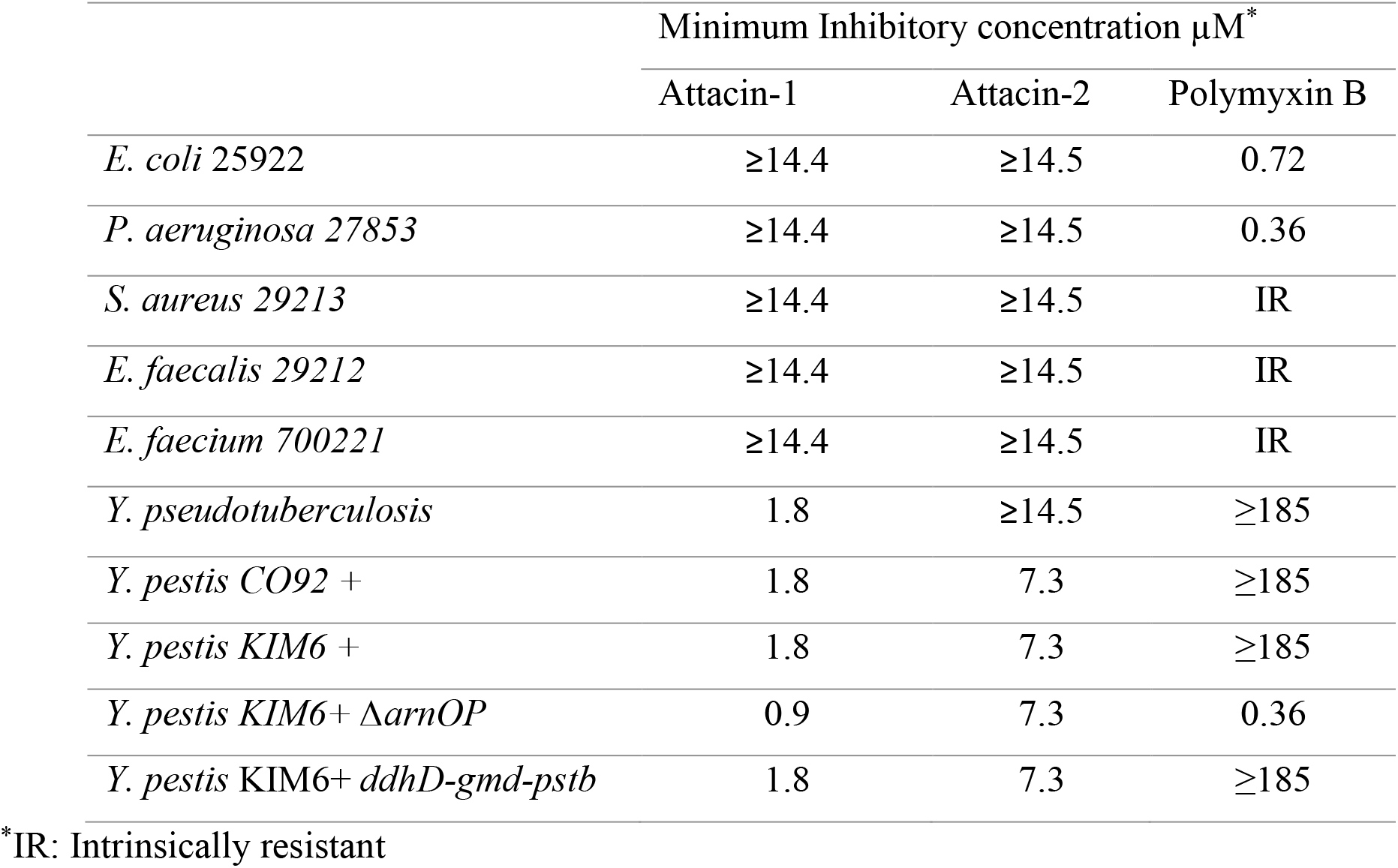
Antibacterial activities of *X. cheopis* attacins.

| | Minimum Inhibitory concentration $\mu\text{M}^*$ | | |
| --- | --- | --- | --- |
|  | Attacin-1 | Attacin-2 | Polymyxin B |
| *E. coli* 25922 | $\geq 14.4$ | $\geq 14.5$ | 0.72 |
| *P. aeruginosa* 27853 | $\geq 14.4$ | $\geq 14.5$ | 0.36 |
| *S. aureus* 29213 | $\geq 14.4$ | $\geq 14.5$ | IR |
| *E. faecalis* 29212 | $\geq 14.4$ | $\geq 14.5$ | IR |
| *E. faecium* 700221 | $\geq 14.4$ | $\geq 14.5$ | IR |
| *Y. pseudotuberculosis* | 1.8 | $\geq 14.5$ | $\geq 185$ |
| *Y. pestis* CO92 + | 1.8 | 7.3 | $\geq 185$ |
| *Y. pestis* KIM6 + | 1.8 | 7.3 | $\geq 185$ |
| *Y. pestis* KIM6+ $\Delta\text{arnOP}$ | 0.9 | 7.3 | 0.36 |
| *Y. pestis* KIM6+ *ddhD-gmd-pstb* | 1.8 | 7.3 | $\geq 185$ |

\*IR: Intrinsically resistant

### Attacins rapidly kill Y. pestis without causing extensive damage to bacterial membranes

To understand the mechanism of action of the attacins, we first examined bacterial killing kinetics (Fig 1A). Treatment with attacin-1 or attacin-2 at 1x MIC concentrations led to a >90% reduction in colony forming units (CFU) of *Y. pestis* in less than 2 h, suggesting rapid bactericidal activity. Unlike the attacin from *Hyalophora cecropia*, which loses its activity in phosphate buffer (*20*), the *X. cheopis* attacins retained activity in PBS (Fig S2). Antibacterial peptides often exert activity by destabilizing bacterial membranes, so we reasoned that the rapid killing exhibited by attacins may be due to permeabilization of *Y. pestis* membranes. Surprisingly, the attacins did not cause substantial damage to *Y. pestis* membranes. Both outer and inner membranes appeared to be largely intact at up to 4x MIC concentration of attacin-1, as there was no observable release of β-lactamase (Fig 1B) nor intracellular accumulation of SYTOX green (Fig 1C). Attacin-2 treatment may lead to slight perturbation of the outer membrane as shown by β-lactamase release, however results were not significantly different from untreated cells (p ≥ 0.084). To further understand the effect on *Y. pestis* KIM6+ membranes, we looked for disruptions in membrane potential due to attacin treatment (Fig 1D). No extensive depolarization was observed, however slight perturbation of the membrane potential was evident with attacin-2. Together, these results indicate that the rapid bactericidal activity of attacins does not involve extensive damage to membranes.

**Fig. 1.**
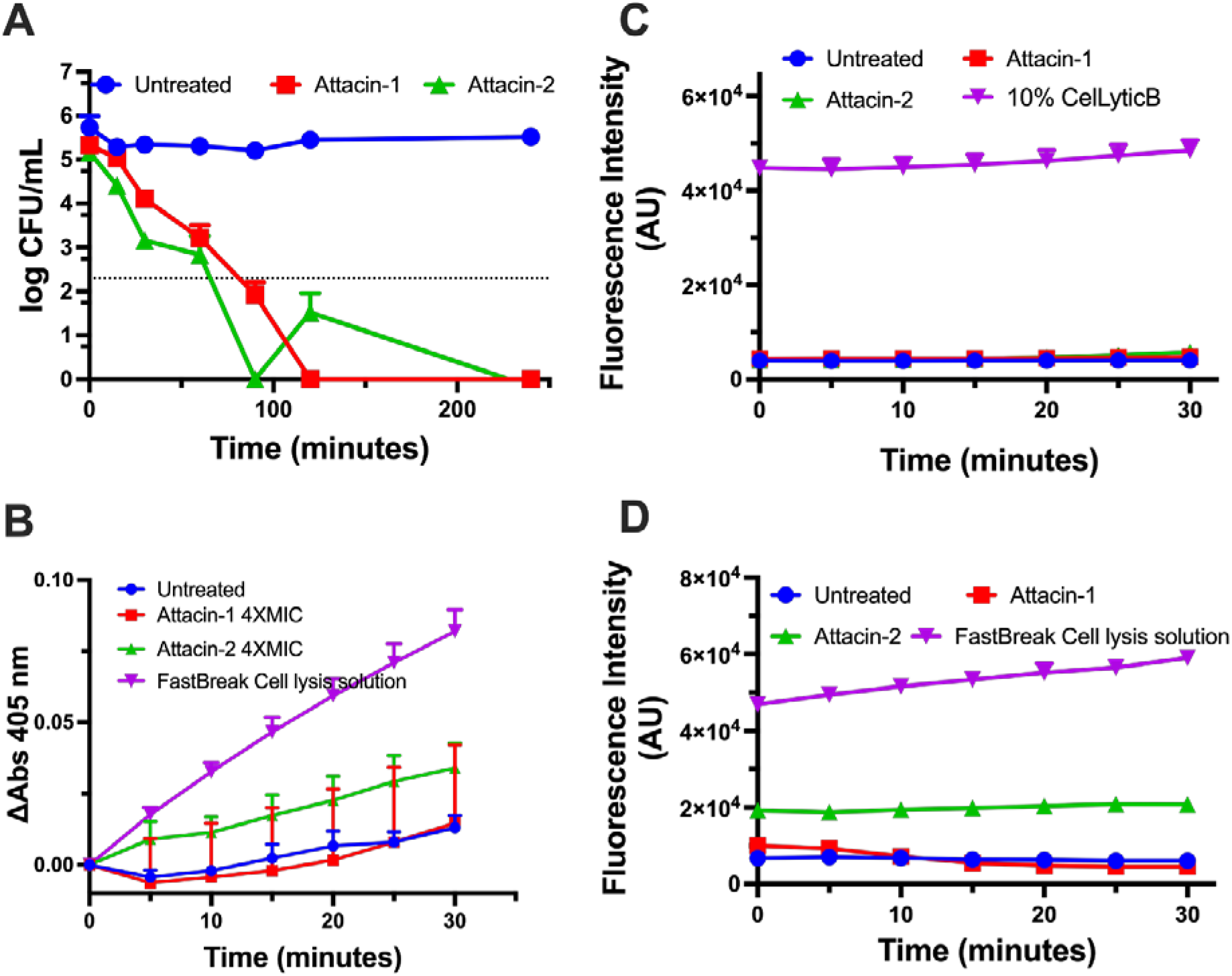
Interactions of *X. cheopis* attacins with *Y. pestis* KIM6+. **(A)** Kinetics of killing of *Y. pestis* KIM6+ by attacin-1 and attacin-2. The dotted lines indicate the limit of detection. **(B)** Effect of attacins on the outer membrane permeability of *Y. pestis* KIM6+ *pUC19*. The extent of permeabilization was quantified by measuring the β-lactamase activity using chromogenic substrate CENTA. **(C)** Effect of attacins on the *Y. pestis* KIM6+ inner membrane. Intracellular accumulation of SYTOX green was monitored by measuring changes in the fluorescence intensities in peptide treated cells. **(D)** Effect of attacins on the membrane potential of *Y. pestis* KIM6+. Changes in the membrane potential were monitored by measuring the fluorescence intensity of the DisC3(5). FastBreak Lysis solution and CelLytic B are detergent controls that permeabilize *Y. pestis* membranes. Values given are the average of 3 independent experiments and error bars represent standard deviation.

### Interactions with LPS

It was previously shown that LPS is the major receptor on *Escherichia coli* for *H. cecropia* attacin (*22, 23*). To assess the ability of *X. cheopis* attacins to interact with LPS, we tested bactericidal activities in the presence of *E. coli* rough LPS. Free LPS inhibited antimicrobial activities of attacins (Fig 2A), with complete inhibition of attacin-2 activity observed at an LPS to peptide ratio [w/w] of 1.25, while attacin-1 still retained 50% activity at the same ratio. Since both attacins have comparable molecular weights (attacin-1: 17844 Da and attacin-2: 17631 Da), it is presumed that attacin-2 possess a higher affinity for *E. coli* LPS. To further assess their affinity for LPS, IC_50_ values were calculated for displacement of dansyl cadaverine (DC) from *E. coli* LPS (Fig 2B). Attacin-2 displaced DC more efficiently with an IC_50_ of 0.6 μM compared to attacin-1 with an IC_50_ of 1.6 μM. Both the bactericidal and DC displacement assays indicated that, like *H. cecropia* attacin, *X. cheopis* attacins are able to bind LPS. Although, attacin-2 has higher affinity for *E. coli* LPS, it showed considerably lower activity against *Y. pestis* compared to attacin-1. The lipid A of *Y. pestis* is modified with Ara4N at 28 °C, and the absence of Ara4N modification substantially enhanced the activity of attacin-2 (Table 1). Therefore, to test the affinities of attacins for LPS modified with Ara4N, we examined their activities in the presence of free *Y. pestis* KIM6+ LPS (Fig 2C). Unlike *E. coli* LPS, *Y. pestis* LPS had minimal effect on the killing activities of attacins. To further test the affinity of attacins for Ara4N-modified vs. unmodified LPS, we carried out whole cell binding assays with fluorescently labelled attacins. The presence of Ara4N reduced the overall affinity of attacins toward *Y. pestis* cells, although the difference was not statistically significant for attacin-1 (p ≥ 0.47, Fig 2D-E). The trend was similar for attacin-2, but a statistically significant reduction in binding due to Ara4N was evident at 4 μM (p = 0.01). Comparable affinities toward wild type and Ara4N mutant cells, and activities in the presence of free KIM6+ LPS indicate potential non-LPS interacting partners on the cell surface.

**Fig. 2.**
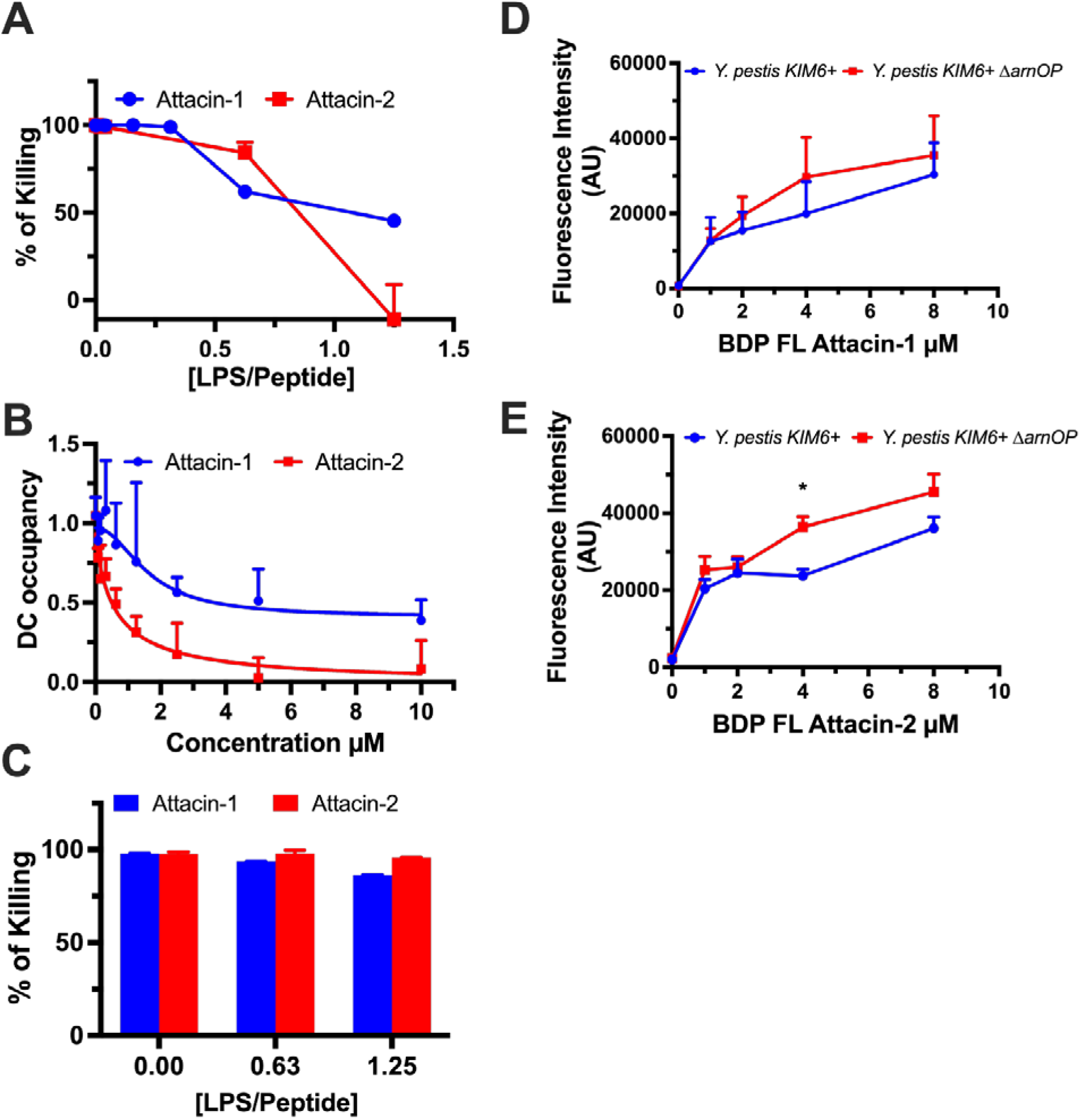
Interactions of *X. cheopis* attacins with LPS. **(A)** Effect of *E. coli* EH100 Ra LPS on the activity of attacins. Bactericidal activity of attacins (at 1X MIC concentration) were tested in the presence of increasing concentrations of LPS. **(B)** The ability of *X. cheopis* attacins to displace *E. coli* EH100 *Ra* LPS bound dansyl cadaverine. **(C)** Effect of *Y. pestis* KIM6+ LPS on the activity of attacins. Purified LPS were incubated with attacins at 1x MIC concentration for 2h and the percentage of killing was calculated. **(D)** Affinity of BDP FL labelled attacin-1 and **(E)** attacin-2 toward *Y. pestis* KIM6+ and *Y. pestis* KIM6+ *ΔarnOP*. Values given are the mean of three independent experiments and error bars represent standard deviations.

### Metabolic activity is essential for attacin-mediated killing

Although the attacins did not rapidly permeabilize bacterial membranes, it is plausible that the interactions with LPS and bacterial outer membranes trigger the generation of reactive oxygen species (ROS) (*25, 26*). Therefore, we quantified the ROS generated in the presence of attacins using the fluorescent probe CM-H2DCFDA (*26*). In the presence of both attacin-1 and -2, rapid generation of ROS was observed in *Y. pestis* (Fig 3A). Interestingly, cells treated with attacin-2 generated considerably higher amounts of ROS compared to attacin-1, despite its higher MIC (Table 1). To further validate the role of ROS in attacin-mediated killing, we analyzed the bacterial killing under anaerobic conditions (Fig 3B) and in the presence of a metabolic inhibitor and protonophore CCCP (Fig 3C). The peptides were able to kill *Y. pestis* under anaerobic conditions, albeit at 10-15% lower rates than in the presence of oxygen, while nearly complete inhibition of killing was observed in the presence of CCCP. To further investigate the role of ROS, we tested the effect of glutathione, a ROS scavenger, on the activity of attacins (Fig 3D). The peptides were able to elicit 90% killing in the presence of this ROS scavenger. The fact that their activity under anaerobic conditions and in the presence of ROS scavengers was moderately reduced suggests that ROS does contribute, albeit to a limited extent, to attacin-mediated killing of *Y. pestis*. Although CCCP inhibits the generation of ROS, it also inhibits broader metabolic activity of the bacteria. Nearly complete survival of *Y. pestis* treated with attacins in the presence of CCCP strongly suggest that metabolic activity plays a crucial role in *X. cheopis* attacin-mediated killing.

**Fig. 3.**
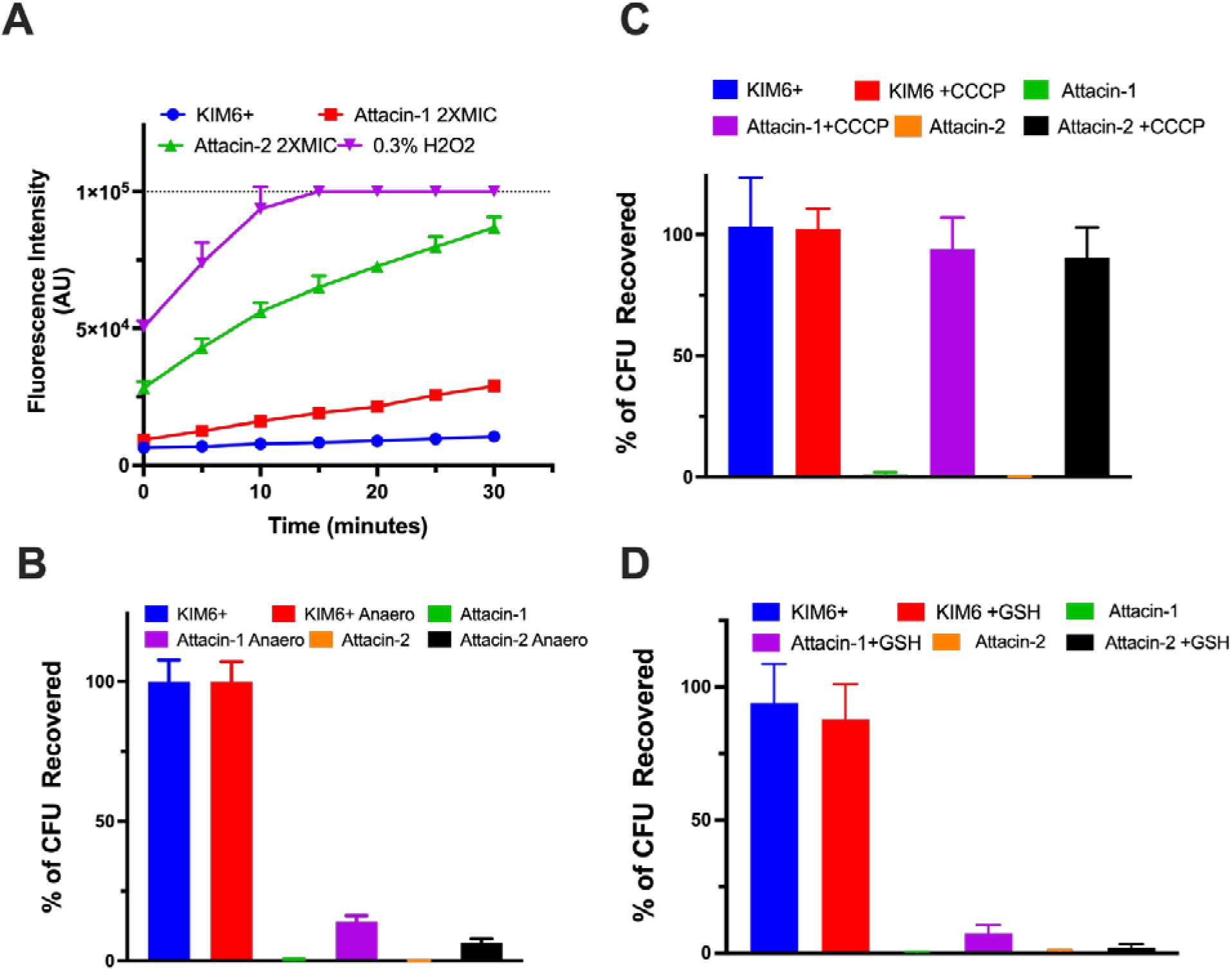
Role of reactive oxygen species and metabolic activity on the activity of attacins. **(A)** Attacin-induced production of ROS in *Y. pestis* KIM6+. The ROS produced was quantified using CM-H2DCFDA by measuring the fluorescence intensity. 0.3% H_2_O_2_ was used as positive control. **(B)** Comparison of bactericidal activities of attacins under aerobic and anaerobic conditions at their respective MIC concentrations. **(C)** Effect of metabolic inhibitor and protonophore CCCP on the activity of attacins and, **(D)** Bactericidal activity of attacins in the presence of ROS scavenger glutathione. Values given are mean and standard deviations of three independent experiments.

### Y. pestis continues to divide in the presence of X. cheopis attacins

Live cell imaging under MIC assay conditions were carried out to get further insights into the mechanism of action of the attacins. Images recorded every 10 min for up to 12 h (Fig 4) revealed the cells continued to grow and divide for several hours in the presence of attacins, though at a reduced rate compared to untreated cells. However, after 6-8 h, a distinct morphological change began to occur, where the bacteria became spherical in shape, followed shortly by lysis. Although both attacin-1 and attacin-2 treated cells showed similar morphological changes, initial changes in morphology were observed around 6 h for attacin-1 while similar changes were evident only at 8 h for attacin-2 treated cells. Further, the cells appeared to grow and divide in the presence of attacin-2 at a relatively higher rate compared to attacin-1 treated cells. The wild type *Y. pestis* KIM6+ are highly resistant to cationic AMPs and detergents, however, we have recently shown that a cecropin class peptide, cheopin, from *X. cheopis* permeabilizes *Y. pestis* in the absence of the Ara4N modification of LPS (*7*). Therefore, we used *Y. pestis* KIM6+ *ΔarnOP*, a mutant lacking Ara4N-modified LPS, to investigate whether the cells undergo similar morphological changes in the presence of the membrane active AMP cheopin. Wild type *Y. pestis* is resistant to cheopin (*7*), and did not show any effect on bacterial growth or morphology. No growth was evident when *Y. pestis* KIM6+ *ΔarnOP* was treated with 1x MIC concentration of cheopin, while attacin treated cells showed similar morphological changes as wild type. However, the attacin-1-associated morphological changes began to occur roughly an hour earlier in the *arnOP* mutant. These results clearly demonstrate that the attacins do not kill *Y. pestis* in a rapidly lytic fashion, with cells continuing to grow for several hours in the presence of the AMPs. It has been previously shown that *H. cecropia* attacin induces filament formation in *E. coli* (*20*). To determine if flea attacins were inducing filament formation in *Y. pestis*, we used confocal microscopy to observe detailed morphological changes induced by attacins (Fig 5). These images clearly indicated the absence of filamentation, and the uniform staining with the membrane dye FM4-64 further suggests that attacins do not induce extensive damage to *Y. pestis* membranes. Interestingly, attacin treatment reduced the total number of cells with GFP fluorescence after 4h, whereas GFP positivity was comparable at 1h (Fig 5). At later times (8h), FM4-64 staining intensity was reduced and high GFP fluorescence was observed outside the cells (Fig S3). These changes are consistent with the rounding and eventual rupture seen in time-lapse imaging of attacin-treated cells.

**Fig. 4.**
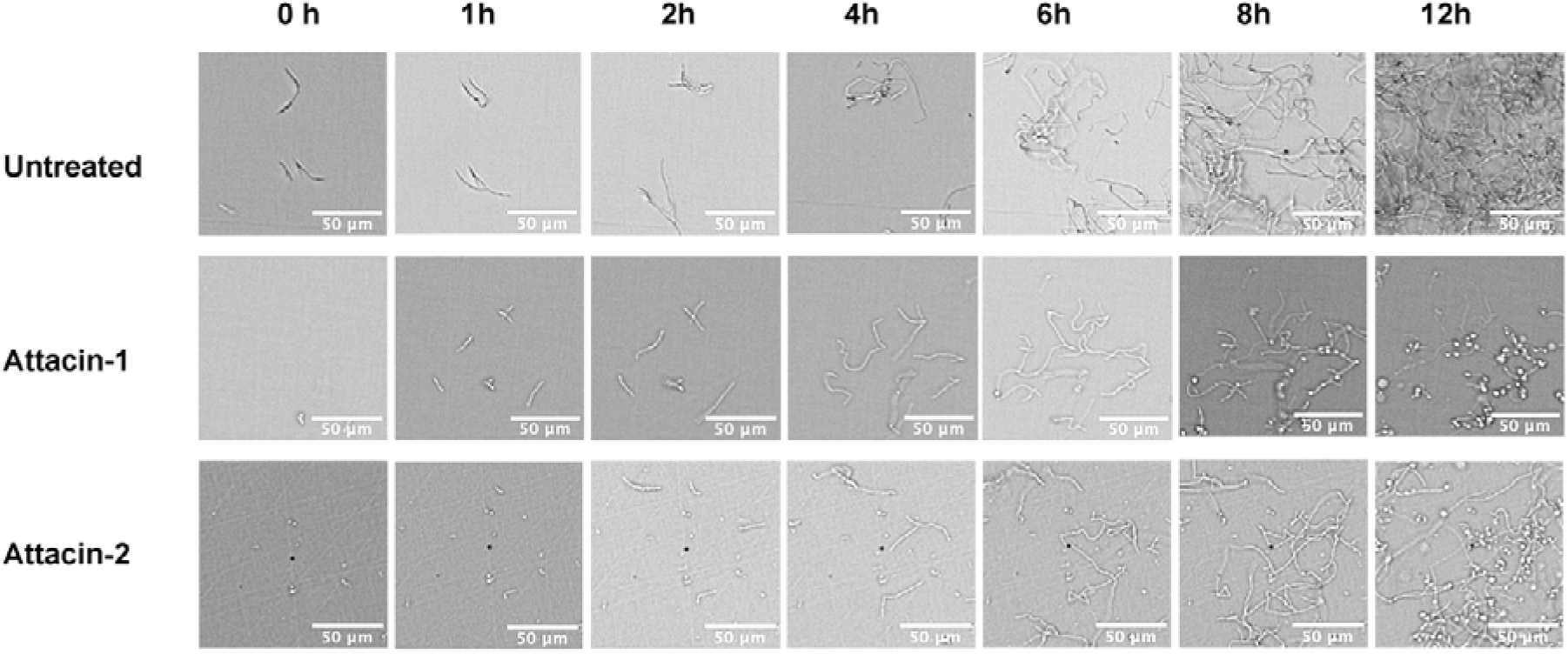
Live cell imaging under MIC-testing conditions. *Y. pestis* KIM6+ cells were treated with MIC concentrations of attacins under MIC testing conditions in cation adjusted Muller-Hinton broth and images were recorded every 10 min for 12h at 28°C.

**Fig. 5.**
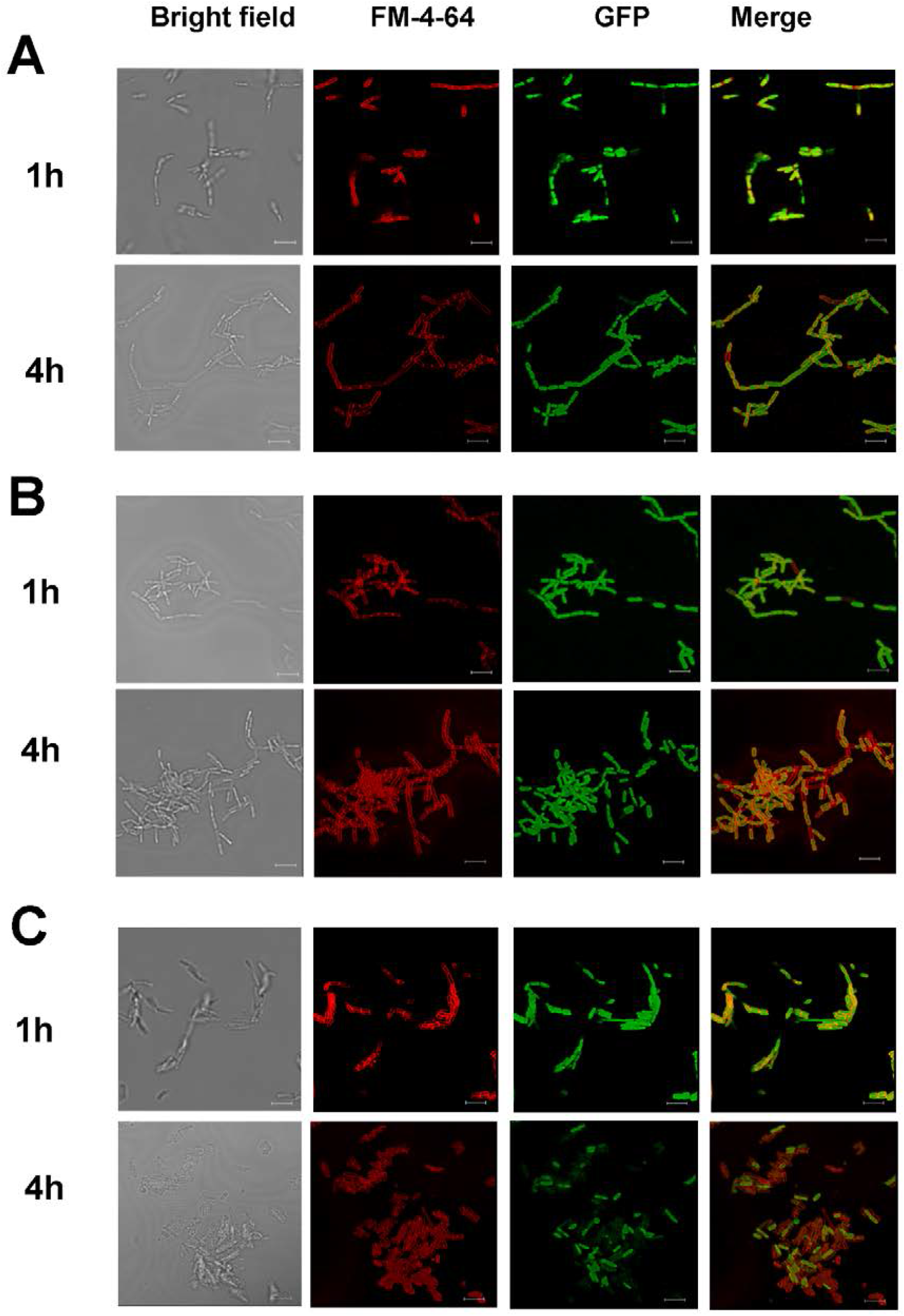
Effect of attacin treatment on *Y. pestis* using high-resolution confocal microscopy. *Y. pestis* KIM6+ expressing GFP was treated with 1X MIC concentrations of attacins for 1h and 4h at 28°C and bacterial membranes were stained with FM4-64 to visualize the cell boundaries. (A) Confocal images of untreated, (B) attacin-1 and (C) attacin-2 treated cells. Scale bar represent 5μm.

## Discussion

In this study, we describe the antimicrobial activities and mechanisms of action of *X. cheopis* attacins. These AMPs showed considerable activity only against *Y. pestis*, and the very closely related *Y. pseudotuberculosis*. Previous studies have shown that insect attacins show considerable activities against certain select Gram-negative bacteria, including bacteria that are resistant to cationic AMPs (*20, 22, 23*). The activity of attacin-1 was somewhat more potent than attacin-2, with a 4-fold lower MIC. Deletion of *arnOP* did not cause any considerable change in the activity of attacin-1 while 8-fold increase in the activity of attacin-2 was observed in the absence of Ara4N-modified LPS. It has been proposed that rough type LPS favors activity of *H. cecropia* attacins (*23*). *Y. pestis* has a rough type LPS (*13*), however, expression of *Y. pseudotuberculosis* O-antigen on *Y. pestis* failed to alter attacin-1 and attacin-2 MICs (Table 1). Further, attacin-1 showed comparable activities against *Y. pseudotuberculosis* and *Y. pestis*, suggesting O-antigen is not a critical factor in determining the activity of *X. cheopis* attacins. *Y. pestis* evolved from *Y. pseudotuberculosis* through several gene gains and losses. However, they share close common ancestry and many common genes and pathways (*27*). The lack of activity against other Gram-negatives, or even other *Enterobacterales*, yet shared activity against these closely-related *Yersinia* species, suggests the *X. cheopis* attacins may have evolved to target a particular aspect of *Yersinia* physiology as a way of minimizing the impact of *Y. pestis* infection.

*H. cecropia* attacins were reported to elicit activity through interaction with LPS and subsequent dysregulation of outer membrane homeostasis, including membrane permeabilization (*22-24*). *X. cheopis* attacins, particularly attacin-2, also bind *E. coli* rough LPS, as shown by displacement of dansyl-cadaverine and significant loss in activity after LPS pretreatment. Intriguingly, unlike *E. coli* LPS, *Y. pestis* LPS did not inhibit the activity of attacins. Further, whole cell binding assays demonstrated comparable affinity for wild type and *arnOP* deleted cells, indicating that Ara4N modification is not primarily responsible for these differences. While LPS may act as a receptor for *X. cheopis* attacins, the lack of substantial disruption of *Y. pestis* membranes suggests a different mechanism of action than has been described for other attacins. It is plausible that other interacting partners on *Yersinia pestis* cells may be responsible for the apparent selectivity and killing activity of these novel attacins.

LPS binding AMPs such as polymyxin B have been reported to induce reactive oxygen species, which contribute to bacterial killing (*26*). Although *X. cheopis* attacins do not rapidly permeabilize *Y. pestis* membranes, considerable amounts of ROS are produced on treatment with these peptides. Under anaerobic conditions and in the presence of ROS scavengers there was a 10-15% inhibition of killing, suggesting that ROS produced in the presence of attacins contribute, albeit marginally, to bactericidal activity. Interestingly, complete inhibition of both attacin-1 and attacin-2 activity was observed in the presence of a metabolic inhibitor CCCP. Further, live cell imaging results showed that cells continue to grow and divide in the presence of attacins. However, after 6-8 h of incubation, a distinct morphological change occurred, followed by cell lysis. The live imaging results of both attacin-1 and 2 were comparable, suggesting both attacins have similar mechanisms of action. Interestingly, killing kinetics assays (Fig 1A) indicated that cells treated with attacins for 2 h do not form colonies when plated in the absence of AMP. Based on the live cell imaging, it is presumable that individual cells continue to grow and divide on solid media, however, they fail to form colonies because they likely rupture after several generations. Unlike the filamentation seen with *E. coli* treated with *H. cecropia* attacin (*20*), confocal images of FM6-64 stained GFP expressing *Y. pestis* cells clearly indicated the absence of filamentation. Further, the leakage of GFP observed at 8h confirms the loss of membrane integrity during the later time points. The fact that *Y. pestis* fails to form colonies after treatment with attacins for 2 h (Fig 1A) suggests that, despite the subsequent limited growth and replication, once exposed to sufficient concentrations of attacin, its fate is sealed. Based on these results, it appears that interactions of attacins with the *Y. pestis* cell surface trigger pathways that lead to energy and/or metabolism-dependent morphological changes and cell death. The physiologic changes induced by the attacins are likely transferred to daughter cells (*28*), as they continue to grow and divide for a limited time in the presence of attacins, yet fail to form colonies when transferred to media lacking attacins.

In conclusion, our results indicate that attacins produced in response to *Y. pestis* infection in fleas exhibit strong and selective activity against *Yersinia*. Although attacins show affinity toward LPS, their mode of action did not involve rapid permeabilization of outer or inner membranes. Instead, they continue to grow and divide for several generations before ultimate disruption. While the reactive oxygen species produced in attacin-treated cells contribute modestly to cell death, the majority of killing occurs via a yet to be described mechanism.

## Experimental Methods

### Cloning, expression and purifications of X. cheopis attacins

The attacin-1 and attacin-2 mRNA sequences reported previously (*10*) were subjected to *in silico* translation (Snap Gene V. 6.0.2, GSL Biotech LLC, Chicago, IL). The protein sequences obtained were processed using Signal P.6 to identify the signal peptide and codon optimized genes of pro-attacins were synthesized for bacterial heterologous expression systems (Genewiz Inc, South Plainfield, NJ). These genes were cloned into pET His6 MBP TEV cloning plasmid (pET His6 MBP TEV LIC cloning vector (2M-T) was a gift from Scott Gradia, Addgene plasmid # 29708), as a maltose binding protein fusion to obtain pETMBP(TEV)Atta-1, and pETMBP(TEV)Atta-2. Cloned peptides had a Tobacco Etch Virus (TEV) protease cleavage site at the N-terminus and digestion with TEV protease added an additional serine residue to the N-terminus of the attacins. Subsequently, pETMBP(TEV)Atta-1, and pETMBP(TEV)Atta-2 were transformed into a *E. coli* BL21DE3 pLys expression strain and peptides were expressed using IPTG (1 mM) induction at 30°C for 4 h. Induced cells were then lysed in the lysis buffer (20 mM Tris, 1 mM EDTA and 200 mM NaCl, pH 8.0) and cell lysate was then subjected to affinity enrichment using Agarose amylose resin [NEB Biolabs Inc, USA], using gravity flow. The resins were then washed with 10X column volume of lysis buffer and amylose-bound proteins were then subjected to on-column TEV protease cleavage in TEV digestion buffer (20 mM Tris, 1 mM EDTA and 200 mM NaCl, 1 mM DTT, and 1 M urea pH 8.0) for 4 h at 30 °C. The TEV protease was expressed and purified in-house using *E. coli* transformed with plasmid pDZ2087 (pDZ2087 was a gift from David Waugh, Addgene plasmid # 92414)(*29*). The cleaved attacins were then eluted from the column and precipitated with 20% ammonium sulfate (*20*). Precipitated peptides were then dialyzed against 25 mM Tris supplied with 1 M urea (pH 8) and subjected to anion exchange chromatography using Pierce strong anion exchange spin columns (ThermoScientific) to remove residual maltose binding protein. The flow through were collected and subjected to ammonium sulfate precipitation, followed by overnight dialysis against 50 mM phosphate buffer (pH 7.5) supplied with 1 M urea. The purity of the final peptides was assessed using SDS PAGE electrophoresis (Fig S1B). The peptides were further characterized by MALDI-TOF mass spectrometry: Attacin-1; 17844.60/17841.62, Attacin-2 17631.15/17630.15 (calculated/observed). Peptides were then quantified using molar absorptivity coefficients 26,930 M^-1^ cm^-1^ and 25,440 M^-1^ cm^-1^ for attacin-1 and attacin-2, respectively. Peptides were flash frozen using a dry ice ethanol mixture and stored at -80°C until further use. The working stocks of the peptides were stored at 4°C, and freezing was avoided, as both attacins aggregated when subjected to refreezing. BDP FL fluorophore-labelled attacins were made using an amine reactive BDP-FL NHS ester (Lumiprobe, Hunt Valley,MD). The fluorophore conjugation was carried out using the manufacturer provided protocol, and as described previously (*7*).

### Antimicrobial activities

The minimum inhibitory concentrations were determined using standardized broth microdilution assays in cation adjusted Mueller Hinton broth (CAMH) as recommended by CLSI (*30*). Assays with *Y. pestis* and *Y. pseudotuberculosis* were carried out at 28°C and the MIC for *Y. pestis* was determined after 36 h (*7*). For all other bacterial strains used in this study, assays were carried out 37°C and MICs were read after 18-24 h of incubation. The minimum bactericidal concentrations (MBC) of the peptides were determined by spreading the cells on AMP-free heart infusion agar (HI) plates at the end of the incubation period. Concentrations at which more than 99.5% killing was observed was taken as the MBC. The killing kinetics assays were performed by plating cells on AMP-free HI plates at 0, 30, 60, 90, 120 and 240 min. The plates were then incubated at 28°C for 36 h and colonies were counted and percentage of killing was calculated.

### Outer membrane permeabilization assays

The effect of attacins on the *Y. pestis* KIM6+ outer membranes were examined by quantifying the β-lactamase activity as described previously (*7*) using the chromogenic substrate CENTA (Sigma). Briefly, *Y. pestis* KIM6+ cells were transformed with pUC19 (*31*) plasmid and cells were selected on an ampicillin containing growth media. Cells from the mid-log phase were then collected and washed in phosphate buffered saline (pH 7.4) supplied with 1 mM CaCl_2_ and 0.5 mM MgCl_2_. The cell density was adjusted to 10^7^ CFU/mL in the same buffer and cells were then treated with 4X MIC concentration of attacins in the presence of CENTA (90 μM) and β-lactamase activity was quantified by measuring the absorbance at 405 nm for 30 min at 28^°^C on a Biotek Synergy H1M plate reader. Fast Break lysis solution (Promega, Madison, WI) was included as positive control.

### Inner membrane permeabilization assay

The extent of cytoplasmic membrane permeabilization by attacins were monitored by quantifying the intracellular accumulation of the membrane-impermeant fluorophore SYTOX green in the presence of attacins (*32*). Briefly, 10^7^ CFU/mL KIM6+ cells were treated with 4X MICs of attacins in the presence of 200 nM SYTOX green in PBS (pH 7.4) and fluorescence intensities were recorded for 30 min at 28^°^C (Ex 490/Em 523 nm) using a Biotek Synergy H1M plate reader. CelLytic B (Sigma) was included as positive control.

### Membrane potential perturbations assay

The effect of attacins on the membrane potential of *Y. pestis* KIM6+ was examined using membrane potential sensitive fluorophore 3,3-dipropylthiadicarbocyanine iodide [DiSC3(5)] (Anaspec Inc, USA) (*33*). Briefly, cells from the mid-log phase cultures were harvested by centrifugation (5000 x g for 5 min at 4^°^C) and washed with 5 mM HEPES buffer (pH 7.4) containing with 5 mM glucose and 0.2 mM EDTA. The cells were resuspended in 5 mM HEPES buffer (pH 7.4) supplemented with 5 mM glucose and cell density was adjusted to 10^7^ CFU/mL in the same buffer. The cells were incubated with 0.5 μM of DiSC3(5) for 15 min at room temperature. DiSC3(5) pretreated cells were subsequently treated with 4X MIC concentrations of attacins and fluorescence was recorded for 30 min at 28^°^C using an excitation wavelength of 622 nm and an emission wavelength of 670 nm on a Biotek Synergy H1M plate reader.

### Effect of LPS on the activity of attacins

The effect of free LPS on the activity of attacins against *Y. pestis* KIM6+ was studied by testing the activity of attacins in the presence of *E. coli* EH 100 rough LPS Ra mutant (Sigma). The MIC concentrations of attacins were preincubated with increasing concentrations of LPS in CAMH. *Y. pestis* KIM6+ cells were then added to this, and final CFU/mL in the assay medium was 2-5 × 10 ^5^ CFU/mL, and incubated for 2 h at 28^°^C. Cells were then plated on a peptide free HI agar, and incubated for 36-48 h at 28^°^C, and colonies formed were counted and percentage of CFU recovered was calculated.

### Dansyl cadaverine and lipopolysaccharide binding assays

The affinities of attacins for *E. coli* rough LPS were assessed by a dansyl cadaverine (DC, Sigma) displacement assay as described elsewhere (*34*), with slight modifications. The *E. coli* EH rough Ra mutant LPS (40 μg/mL) was incubated with 100 μM DC in 2 mM Tris buffer pH 7.2 for 5 min at room temperature. This mixture was then treated with increasing concentration of peptides for 5 min at room temperature and fluorescence was quantified using excitation and emission wavelengths, 340 nm and 500 nm respectively. The DC occupancy was calculated using the formula: *DC Occupancy*=(*F*- *F*_0_)(*F*_*max*_-*F*_0_), where *F*_*max*_ is the fluorescence of DC the in the presence of LPS, *F*_*0*_ is the fluorescence of DC in the buffer without LPS and, *F* is the fluorescence of DC the in the presence of LPS when treated with peptide.

### Whole cell binding assays

The affinities for *X. cheopis* attacins for *Y. pestis* KIM6+ and *Y. pestis* KIM6+ *ΔarnOP* were quantified using a whole cell binding assay, as described previously (*7*). Cells were incubated with BDP FL labelled attacins and the fluorescence of the cell-bound peptides were quantified by measuring fluorescence (Ex 490/Em 520) on a Biotek Synergy plate reader.

### Reactive oxygen species measurements

The reactive oxygen species (ROS) produced by *Y. pestis* KIM6 + in the presence of attacins were measured using the fluorescent probe CM-H2DCFDA (ThermoFisher) (*26*). Briefly, cells growing from the mid log phase were collected by centrifugation (5000 x g, 5 min), washed and resuspended in PBS. Cells were incubated with 2 μM CM-H2DCFD for 5 min at room temperature. The cells were then treated with 2X MIC concentration of attacins and fluorescence intensities were measured on a Biotek Synergy plate reader using excitation and emission wavelengths of 490 nm and 523 nm, respectively.

### Effect of ROS scavenger glutathione on the activity of attacins

The effect of ROS scavengers on the bactericidal activity of attacins was tested by co-incubating attacins and glutathione (GSH), a known ROS scavenger (*35*). *Y. pestis* KIM6+ cells were treated with 1X MIC concentration of attacins in the presence of 1 mM GSH for 2h. Cells were then transferred to AMP-free HI plates and incubated for 36-48 h at 28^°^C. Colonies were counted and percentage of killing was calculated.

### Effect of CCCP on the activity of attacins

Effect of the protonophore and metabolic inhibitor carbonyl cyanide 3-chlorophenylhydrazone (CCCP, Sigma) on the activity of attacins was measured by co-incubating 50 μM CCCP and 1X MIC concentrations of attacins, as described for LPS and ROS scavenger effects. After 2h of incubation, cells were transferred to AMP-free HI plates and incubated for 36-48 h at 28^°^C. Colonies were counted and percentage relative to untreated CFUs were calculated.

### Live cell imaging of Y. pestis KIM6+ under MIC conditions

The in-solution live images of the *Y. pestis* KIM6+ and *Y. pestis* KIM6+ *ΔarnOP* were recorded using an oCelloscope instrument (BioSense Solutions ApS, Denmark) under MIC testing conditions. Briefly, cells were incubated with or without 1X MIC concentration of attacins or 8 μg/mL of cheopin in CAMH broth at 28^°^C in a flat bottom polystyrene plate (Greiner bio-one, USA). The images were recorded every 10 min for up to 12 h. The recorded images were processed using UniExplorer software (BioSense Solutions ApS, Denmark) and saved as still images or time lapse videos (8 frames/sec).

### Confocal microscopy studies

*Y. pestis* KIM6+ cells (100 μL, 10^7^ CFU/mL) expressing GFP were treated with 1X MIC concentration of attacins in CAMH at 28°C for 1, 4 and 8 h. At the end of incubation 1 μM of FM4-64FX (ThermoFisher) was added to the cells to stain the bacterial membranes and left at room temperature for 5 min. 10 μL of this FM 4-64FX stained cells were then transferred to low melting agarose pads and cells were imaged using a Leica SP8 Lightning confocal microscopy imaging system using a 63x oil immersion objective (NA 1.4). GFP and FM4-64FX were excited using argon 488 nm and HeNe 561 nm laser lines, respectively. The images collected were processed for brightness and contrast using LAX software (Leica Microsystems).

### Statistical Analysis

All experiments were conducted independently at least 3 times and values given are the mean and standard deviations. Unpaired t-tests were carried out to find the statistical significance (p ≤ 0.05). All the graphs plotted and analysis done are using Graph Pad prism. The microscopy images given are representative images or videos from independent experiments.

## Acknowledgments

We thank the HPLC and Imaging core facilities of University of Utah for their help with HPLC and confocal Imaging. We would also like to thank BioSense Solutions ApS, Denmark kindly providing the oCelloscope to carry out live imaging.

## Author contributions

Conceptualization: BM, KLA MF

Methodology: BM, KLA

Investigation: BM, KLA

Visualization: BM

Supervision: MF

Writing—original draft: BM, MF

Writing—review & editing: BM, KLA, MF

