## Supplemental Information for "Antibacterial activity of *Xenopsylla cheopis* attacins against *Yersinia pestis*"

Basil Mathew[1], Kari L Aoygi[1] and Mark A. Fisher*[1,2]

[1]University of Utah Department of Pathology, [2]ARUP Laboratories, Salt Lake City, UT, 84112, United States of America

**This PDF file includes:**

Supplementary Text
Figs. S1 to S3
Tables S1 and S2
References

### Supplementary Text

#### Supplementary Methods:

**Construction of *Y. pestis* KIM6+*::PcysZK-gfp*G**: A stable green fluorescent protein (GFP) expressing *Y. pestis* KIM6+ was constructed using the method described previously [1]. Briefly, the gene encoding GFP under a *PcysZK* promoter was cloned into the mini-Tn7 vector pUC18R6KT-mini-Tn7T-Km [2], which allows innocuous chromosomal insertion into the Tn7 insertion site in *Y. pestis* using the helper plasmid pTNS2. Cells were then selected for kanamycin resistance and recovered clones screened for *gfp* using PCR and visualization. The kanamycin resistance marker was subsequently removed using Flp-recombinase mediated excision using plasmid pFLP2.

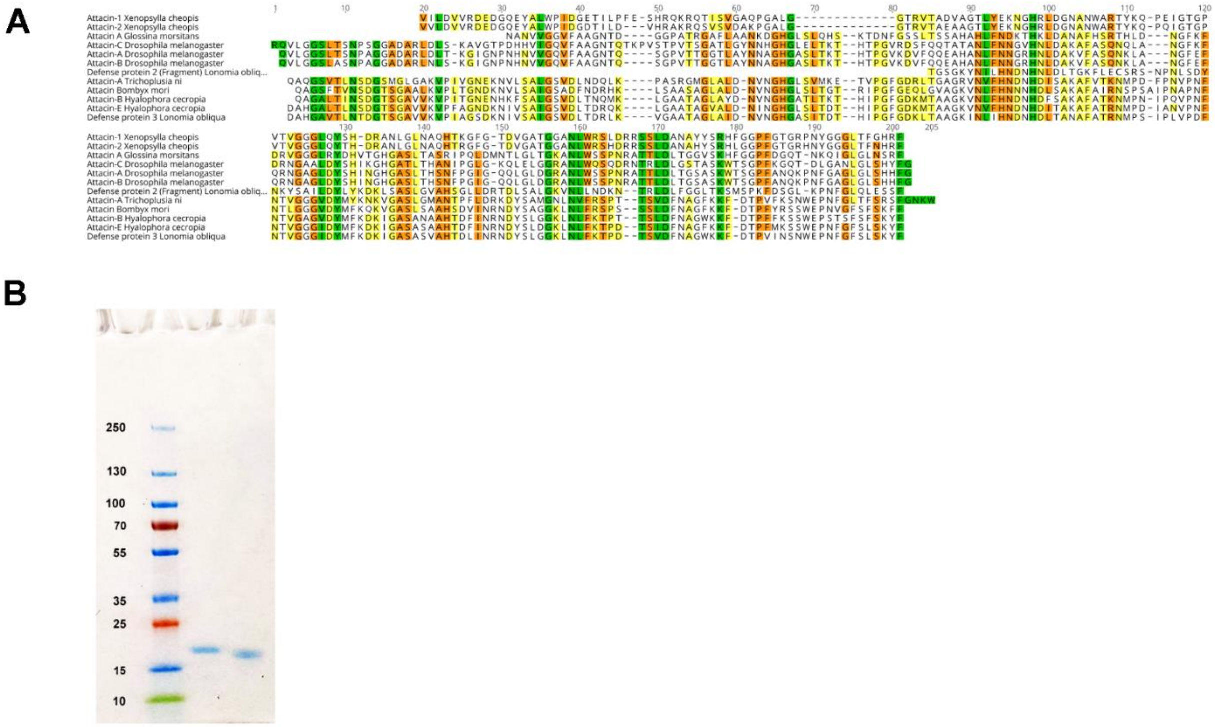


**Fig. S1.** ***X. cheopis*** **attacins cloning and expression.** (A) Amino acid sequences of *X. cheopis* attacins aligned against other known attacin sequences. (B) SDS-PAGE gel images of purified attacin-1 (lane 2) and attacin-2 (lane 3). Lane 1 contains pre-stained protein ladder; numbers indicate molecular weights in kDa.

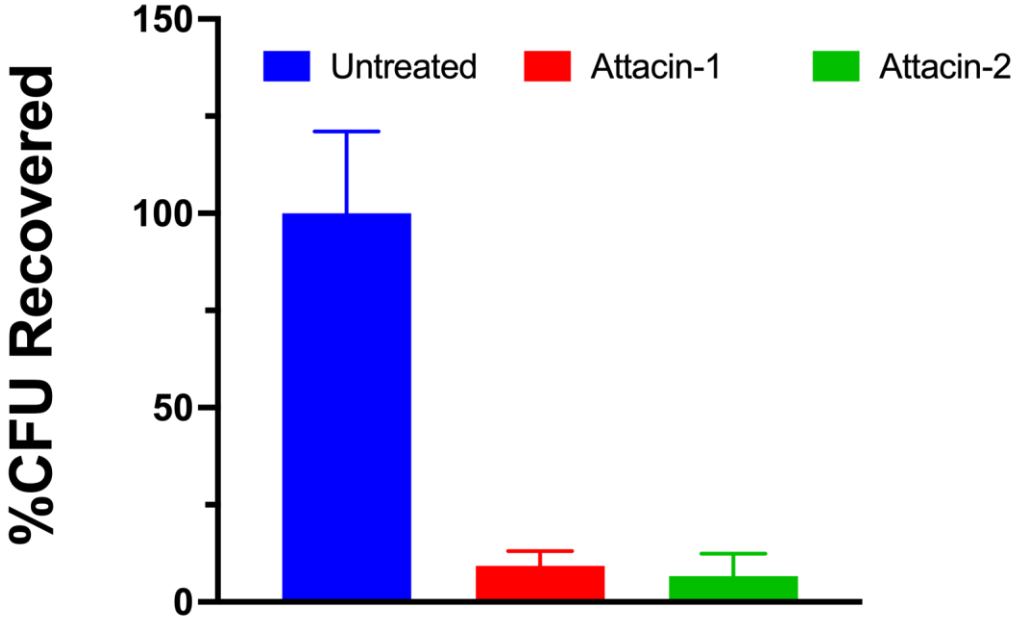


**Fig. S2. Antibacterial activity of attacins.** *Y. pestis* KIM6+ cells were treated with 1X MIC concentration of attacins in PBS (pH 7.4) for 2h at 28 °C and the percentage of killing calculated. Values are the average of three independent experiments and error bars represent standard deviations.

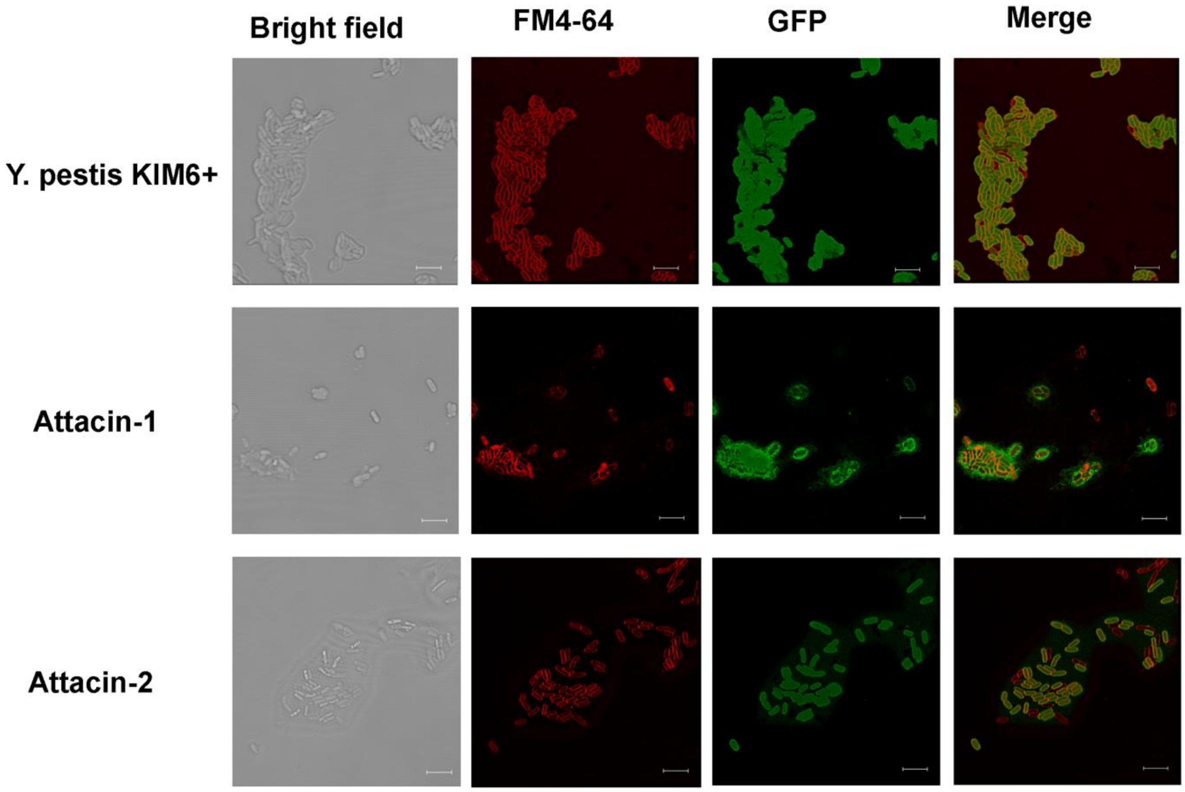


**Fig S3. Confocal microscopy images of *Y. pestis* KIM6+ treated with attacins for 8h.** *Y. pestis* KIM6+ cells expressing GFP were treated with MIC concentrations of attacins for 8h at 28 °C, membranes were stained with FM4-64 and images were recorded. Scale bar represent 5 µm.

**Table S1:** Antibacterial activity of attacins against polymyxin B resistant bacteria. MICs are shown in µM.

| | Attacin-1 | Attacin-2 | Polymyxin B |
|---|---|---|---|
| *Proteus mirabilis* 10195 | ≥14.4 | ≥14.5 | ≥185 |
| *Serratia marcescens* ATCC 8100 | ≥14.4 | ≥14.5 | ≥185 |
| *E. coli* AR#0493 (*mcr-1*) | ≥14.4 | ≥14.5 | 2.9 |
| *Acinetobacter baumannii* complex ci1 | ≥14.4 | ≥14.5 | 2.9 |
| *A. baumannii* complex ci2 | ≥14.4 | ≥14.5 | 1.4 |
| *A. baumannni* complex ci3 | ≥14.4 | ≥14.5 | 46.3 |

**Table S2.** Bacterial strains used in this study.

| Strain | Genotype/ Description | Source/Reference |
|---|---|---|
| *S. aureus 29213* | Standard strain for CLSI antimicrobial susceptibility testing | ATCC |
| *E. faecalis 29212* | Standard strain for CLSI antimicrobial susceptibility testing | ATCC |
| *E. feacium700221* | Standard strain for CLSI antimicrobial susceptibility testing | ATCC |
| *E. coli 25922* | Standard strain for CLSI antimicrobial susceptibility testing | ATCC |
| *P. aeruginosa 27853* | Standard strain for CLSI antimicrobial susceptibility testing | ATCC |
| *Y. pestis CO92* | parental *Y. pestis* strain; pCD−, pMT+, pPCP+ | [3] |
| *Y. pestis* KIM6+ | parental *Y. pestis* strain; pCD−, pMT+, pPCP+ | [4] |
| *Y. pestis* KIM6+ Δ*arnOP* | In frame, markerless deletion of *arn* operon *(arnBCADTEF)* in KIM6+ | [5] |
| *Y. pestis* KIM6+ *dhD-gmd-pstb* | O-antigen pseudogene locus in *Y. pestis* KIM6+ is replaced with functional O-antigen locus from *Y. pseudotuberculosis* | [6] |
| *Y. pestis* KIM6+ pUC19 | *Y. pestis* KIM6+ harboring pUC19 plasmid for β-lactamase expression | [5] |
| *Y. pestis* KIM6+::*PcysZK-gfp* | *Y. pestis* KIM6+ that harbor constitutively expressing GFP inserted into the chromosome | This study |
| *Y. pseudotuberculosis* | Clinical isolate | ARUP |
| *P. mirabilis*10195 | Wildtype *P. mirabilis* | Gift from Dr. Brian Luna, University of Southern California. |
| *S. marcescens* 8100 | Standard strain for CLSI antimicrobial susceptibility testing | ATCC |
| *E. coli* AR#0493 mcr-1 | *E. coli* harboring *mcr-1* plasmid for colistin resistance | CDC |
| *A. baumannii complex ci1* | Colistin-resistant clinical Isolate | ARUP |
| *A. baumannni complex ci2* | Colistin-resistant clinical Isolate | ARUP |
| *A. baumannii complex ci3* | Clinical Isolate | ARUP |
